# Dendrimer-targeted immunosuppression of microglia reactivity super-accelerates photoreceptor regeneration kinetics in the zebrafish retina

**DOI:** 10.1101/2020.08.05.238352

**Authors:** Kevin B. Emmerich, David T. White, Siva P. Kambhampati, Grace Y. Lee, Tian-Ming Fu, Arpan Sahoo, Meera T. Saxena, Eric Betzig, Rangaramanujam M. Kannan, Jeff S. Mumm

## Abstract

Müller glia (MG) function as injury-induced retinal stem cells in zebrafish but not mammals. Insights from zebrafish, however, have been used to stimulate limited regenerative responses from mammalian MG. Microglia/macrophages regulate MG stem cell activity in the chick, zebrafish and mouse. We previously showed that dexamethasone can accelerate retinal regeneration in zebrafish. Similarly, microglia ablation enhances regenerative outcomes in the mouse retina. Targeted immunomodulation may therefore enhance the regenerative potential of human MG. Nanoparticle-based immunomodulation is an emerging field with immense therapeutic potential. Here, we investigated how regeneration-enhancing dexamethasone treatments alter microglia behavior and how dendrimer-based targeting of dexamethasone to reactive microglia impact retinal regeneration kinetics. Intravital time-lapse imaging revealed specific dexamethasone-induced changes in microglia reactivity. Dendrimer-conjugated dexamethasone treatments resulted in: 1) decreased toxicity, 2) selective targeting of reactive microglia and, 3) “super-accelerated” retinal regeneration kinetics. These data support the use of dendrimer-based drug formulations for modulating microglia reactivity in degenerative disease contexts, especially as therapeutic strategies for promoting regenerative responses to neuronal cell loss.

## Introduction

Neurodegenerative diseases are caused by the loss of discrete neuronal cell types. Therapies for regenerating lost neurons are therefore needed to restore neural function. Studies in zebrafish, a naturally regenerative species, have revealed factors that can stimulate neural repair in mammalian models. For instance, retinal Müller glia (MG), which function as injury-inducible neural stem cells in zebrafish (1, 2), can be stimulated to produce new neurons in mammals (3–5). Enhancing the regenerative capacity of MG has the potential for immense therapeutic impact by enabling the restoration of visual function (6).

Studies in the chick initially implicated microglia/macrophages as regulators of MG reactivity and proliferation in response to neuronal cell loss (7, 8). Further, we showed that immunosuppression can accelerate rod photoreceptor regeneration in the zebrafish retina (9). More recently, microglia ablation was shown to enhance regeneration in the mouse retina (10). Importantly, in fish, the timing of immunosuppression profoundly altered the effect on regeneration. Exposure to the glucocorticoid dexamethasone (Dex) prior to retinal cell loss inhibited the regenerative process. However, when applied after the onset of cell loss, Dex accelerated rod cell replacement (9). Similarly, acute inflammatory responses promote regeneration of the optic nerve in zebrafish, but prolonged Dex treatments inhibited repair (11). Together, these results suggest context-dependent modulation of immune system reactivity could be used to promote regenerative responses to neuronal loss.

Unfortunately, systemic immunomodulation incurs deleterious side effects. Nanomedicine-based targeted drug delivery to reactive immune cells can overcome this issue by reducing systemic toxicity and enhancing therapeutic efficacy. For example, we have shown that hydroxyl (poly)amidoamine (PAMAM) dendrimer nanoparticles (12, 13) enable targeting of drug conjugates to reactive microglia/macrophages in multiple disease models (14, 15). In particular, dendrimer-based targeting of Dex to activated macrophages led to improved clinically-relevant measures in a rabbit model of autoimmune dacryoadenitis (16). These data demonstrate that dendrimer-drug conjugates can quell neuroinflammation in neurodegenerative contexts (14). To expand these studies to neuroregenerative contexts, we investigated how dendrimer-based targeting of Dex to reactive macrophages/microglia impacts rod photoreceptor regeneration kinetics (9).

Using a selective rod photoreceptor ablation model (9, 17), we first investigated the effects of post-ablation Dex treatments on microglia reactivity in the degenerating retina. Adaptive optical lattice light-sheet microscopy (AO-LLSM) based *in vivo* time-lapse imaging (18), was used to follow microglia dynamics at high spatial and temporal resolution following induction of rod cell loss. Intravital time-series confocal imaging of dendrimers conjugated to fluorescent reporters also allowed us to assess the dynamics and specificity of cellular targeting at the whole-organism level. Finally, comparisons between free Dex and dendrimer-conjugated Dex (D-Dex) treatments were made to assess relative effects on toxicity and rod cell regeneration kinetics. The data show that post-ablation Dex treatments suppress microglial reactivity to rod cell loss. Moreover, relative to free Dex controls, D-Dex conjugates reduced toxicity, promoted selective targeting of dying neurons and reactive immune cells, and “super-accelerated” rod cell regeneration kinetics. These findings support the development of nanotechnology-based immunomodulation strategies for promoting retinal repair.

## Results

### Post-ablation Dex treatment suppresses microglia reactivity to rod cell death

Our initial characterization of the effects of immunosuppression on retinal regeneration indicated that Dex could either inhibit or promote rod photoreceptor regeneration depending on the timing of treatment. Pre-treating with Dex, prior to induction of rod cell ablation, inhibited regeneration. However, when Dex was applied a day after rod cell loss had been initiated, rod cell regeneration kinetics accelerated (9). Mechanistically, *in vivo* time-lapse imaging showed that pre-treating with Dex inhibited microglia reactivity to rod cell death. However, how microglia dynamics were affected by post-ablation Dex treatment, the paradigm that enhanced regeneration, remain unknown. To address this, we used a powerful intravital microscopy technique developed by the Betzig lab, AO-LLSM (18, 19) to increase spatial and temporal resolution of microglia/macrophage dynamics during *in vivo* time-lapse imaging of larval zebrafish retinas. Specifically, 30-90 micron retinal volumes were scanned at time scales ranging from 5 to 180 sec/intervals – i.e. as much as 120 times faster than our prior study – to assess the effects of post-ablation Dex treatment on microglia/macrophage behaviors near dying rod cells.

For AO-LLSM, a transgenic line enabling selective ablation of fluorescently labeled rod photoreceptors (i.e., rod cells expressing a YFP-NTR fusion protein) was crossed with a transgenic line labeling microglia/macrophages with a complementary fluorescent reporter (tdTomato). Dual-labeled transgenic larvae were selected and treated with the prodrug metronidazole (Mtz) at 5 days post-fertilization (dpf) to induce ablation of YFP-NTR expressing rod photoreceptors (with non-ablated “No Mtz” larvae serving as controls). After 24 h, Mtz was removed and larvae in each group, ablated and non-ablated, were treated with either 2.5 μM Dex or control media. AO-LLSM was then used for high-resolution time-lapse imaging of interactions between microglia and rod cells in all four treatment groups starting at 6.5 dpf (12 h post-Dex treatment). After imaging, labeled cells were 3D-rendered in Imaris to facilitate automated quantification of migration speed and sphericity of microglia/macrophage as metrics of reactivity. Microglia in non-ablated control fish, treated with or without Dex (±Dex), typically exhibited ramified morphologies, actively extending and retracting cellular processes, but remained relatively non-migratory, consistent with a non-reactive homeostatic state (Fig. 1A, B, Supp. Videos 1 and 2). Conversely, microglia in the ablated groups had largely spheroid morphologies suggestive of reactivity. Microglia in Mtz-treated larvae that were not treated with Dex became highly motile, rapidly migrating in and among the dying rod cells (Fig. 1C, Supp. Video 3). Conversely, despite retaining amoeboid morphologies, microglia in ablated larvae treated with Dex remained static (Fig. 1D, Supp. Video 4). Quantitative analysis of sphericity suggested a transition from ramified to amoeboid profiles among microglia in ablated larvae (Fig. 1E). Quantification of migration speed showed no statistically significant differences between non-ablated controls (±Dex) or ablated larvae treated with Dex. Conversely, microglia in ablated larvae that were not treated with Dex were 27-37% more motile than all other groups, a statistically significant difference in all cases (Fig. 1F). Notably, the enhanced temporal and spatial resolution afforded by AO-LLSM allowed us to observe highly motile peripheral macrophages entering the retina in response to rod cell ablation (Supp. Fig. 1 and Supp. Videos 5 and 6). This data clarifies that the innate immune response to rod cell loss in this model system is not limited to tissue-resident microglia, as suggested by our prior time-lapse imaging experiments at much lower temporal resolution (9), highlighting the value of AO-LLSM for accurately monitoring highly dynamic cellular behaviors.

**Figure 1.**
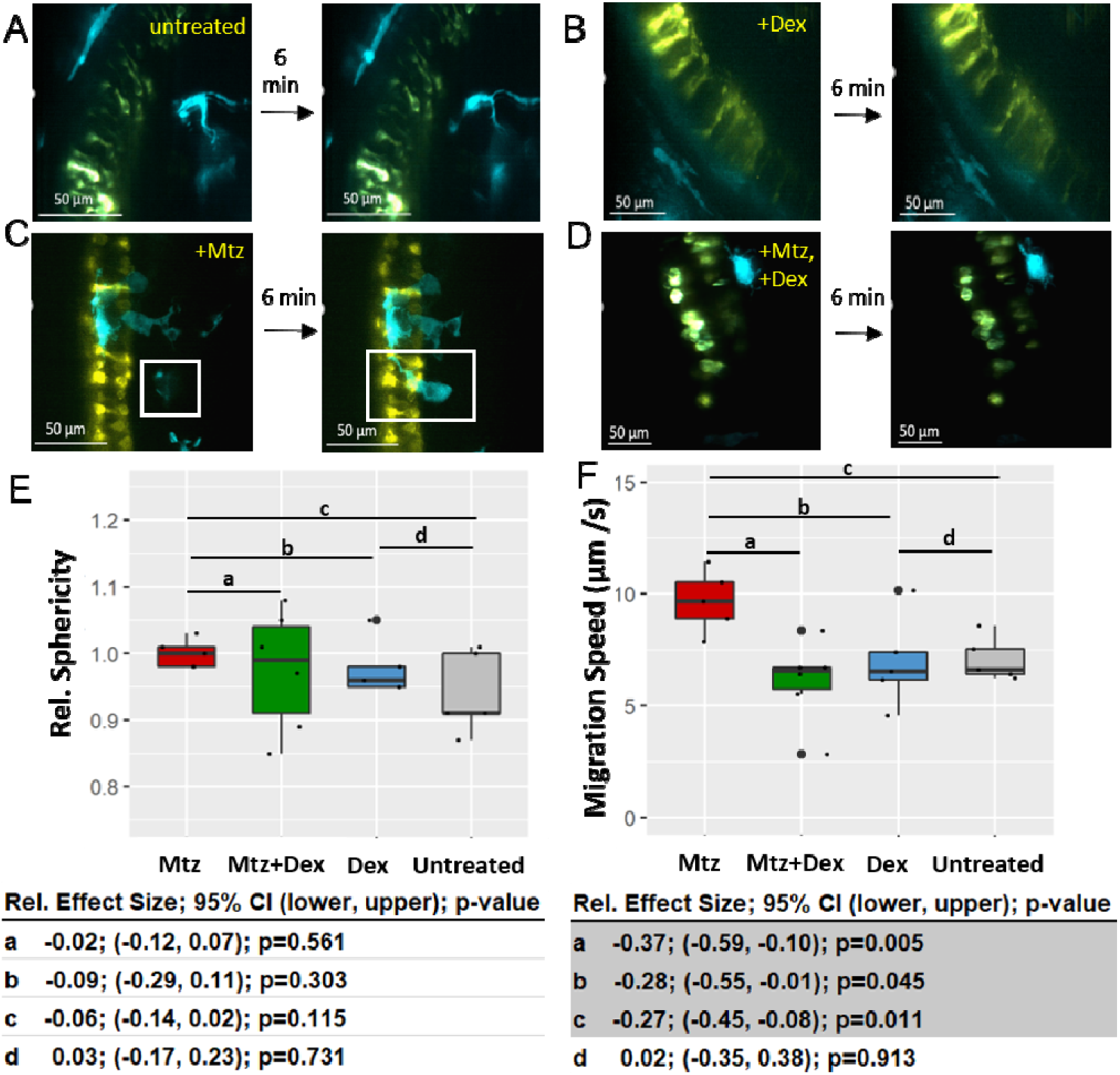
Microglia migration speed is suppressed in response to post-ablation Dex treatment. **A-D**) Image stills from AO-LLSM time-lapse imaging (30 sec time intervals) of rod photoreceptors (yellow, NTR-YFP) and microglia/macrophages (cyan, false coloring of tdTomato). Four conditions were imaged including non-ablated control (**A**; untreated), non-ablated Dex-treated (**B**; +Dex), rod cells ablated (**C**; +Mtz) and rod cells ablated plus Dex (**D**; +Mtz, +Dex). Fish were imaged at 5 or 6 dpf using AO-LLSM. **A-B**, Microglia in untreated fish and fish that received Dex only were ramified and often extended and contracted processes but exhibited little to no migration. **C**, Microglia in fish whose rod cells were ablated with Mtz (10 mM, 12 or 24 hours) migrated rapidly to and interacted with dying rod cells visible (white box shows one microglia entering the frame and then interacting with dying rods). **D**, Fish soaked with Dex post-Mtz (2.5 μM, 6 or 24 hours) exhibited comparably static microglia that did not migrate toward dying rod cells. **E-F**, Imaris was used to quantify relative sphericity and migration speed (μm/second) of microglia in each group (6-8 fish per condition). Statistical comparisons between groups (a-d) were done to calculate effect size, 95% confidence intervals, and p-value. A significant difference was observed between the speed of microglia treated with Mtz+Dex compared to Mtz only (−37%, **F**) while a minimal difference was observed in relative sphericity (−2%, **E**). See supplemental videos 1-4 corresponding to stills A-D, respectively.

### Dendrimer-conjugated dexamethasone (D-Dex) shows little to no toxicity

Long-term therapeutic use of Dex and other glucocorticoids is associated with complications due to multiple adverse side-effects and systemic toxicity concerns (20). Therefore, we were interested in exploring whether conjugating Dex to dendrimers could reduce deleterious side-effects, as well as target Dex to reactive microglia, as per prior studies (15, 16, 21). For this, we conjugated a generation 4 (G4-OH) PAMAM dendrimer to Dex (D-Dex). D-Dex conjugates used in this study were >98% pure and have ~8 molecules of Dex conjugated to the surface hydroxyl groups of the PAMAM dendrimers, as reported previously (22). To test toxicity, zebrafish larvae were injected in the pericardium (PC) with Dex, D-Dex or vehicle alone at 5 days-post fertilization (dpf) at concentrations that ranged from 2.5 to 400 μM (Fig. 2A; note, injections were necessary as soaking in D-Dex showed no evidence of bioactivity). Mortality rates for each condition were measured two days following injection (7 dpf).

**Figure 2.**
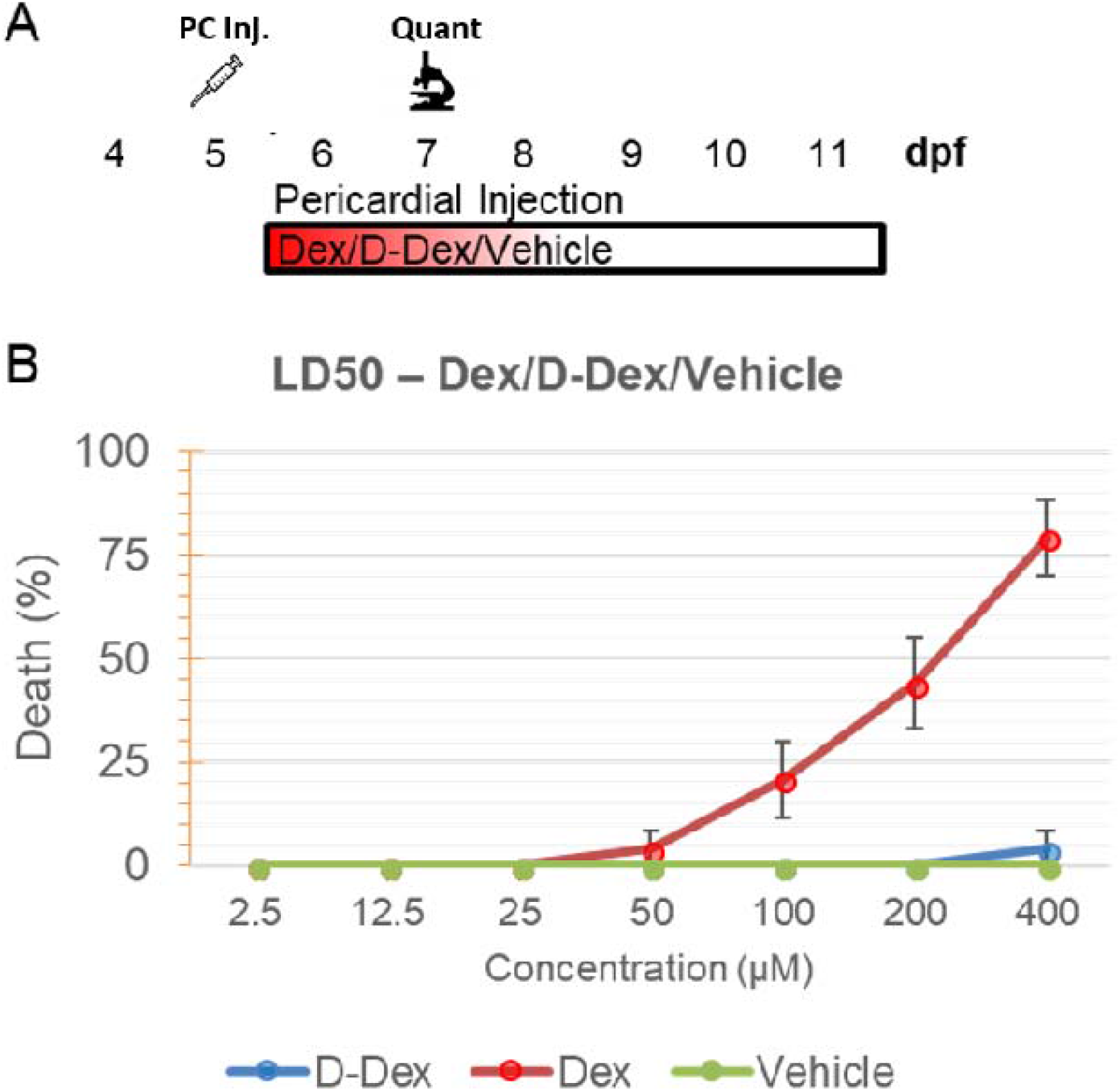
D-Dex injection has reduced toxicity in larval zebrafish. **A,** Schematic of injection assay to test D-Dex toxicity. At 5 dpf, D-Dex, Free-Dex (Dex), or control vehicle were injected into the pericardium of zebrafish larvae (≤10nL injected volume, 2.5-400 μM for D-Dex or Free-Dex). At 7 dpf, toxicity was quantified based on percent survival. **B**, Line graph demonstrating percent of dead fish at 7 dpf injected with vehicle (green line), Dex (red line) or D-Dex (blue line, concentrations indicated along x-axis). Fish injected with Dex displayed the greatest toxicity (LD50 estimated ~207 μM), with fish beginning to die at 50 μM and higher concentrations. D-Dex had zero toxicity at ≤200μM, and minimal toxicity at 400 μM, closely matching the vehicle control.

At lower concentrations, 2.5 to 25 μM, no death was observed amongst the three injection groups (Dex, D-Dex or vehicle). However, Dex-injected fish had increased mortality rates at higher concentrations, reaching 75% toxicity at 400 μM (LD_50_ ~ 207 μM, Fig. 2B). In contrast, no toxicity was observed in the vehicle injected group and only a ~5% toxicity rate was seen in the 400 μM D-Dex injected group (Fig. 2B). These findings are in agreement with previous studies in which D-Dex attenuated Dex toxicity ~10-fold (22).

Next, to investigate the pharmacokinetics of dendrimer compounds in zebrafish we injected FITC-labeled dendrimers (D-FITC) into the PC of 5 dpf healthy zebrafish expressing *Tg(mfap4.1:Tomato-CAAX)*. Subsequently, fish were subject to confocal microscopy at 1 and 2 days post injection (dpi). The volume of D-FITC fluorescent signal in the zebrafish kidney was quantified using Imaris. D-FITC in each fish at 2 dpi was compared to the same fish at 1 dpi to quantify the pharmacological clearing of dendrimers. By 1 dpi, the majority of D-FITC accumulates in the kidney. On average, 27% of D-FITC was cleared between 1 and 2 dpi demonstrating rapid clearing of injected dendrimers in larval zebrafish (Supp. Fig. 2 A, B).

### D-Dex post-ablation treatment super-accelerates rod photoreceptor regeneration

We previously observed a ~30% increase in regeneration kinetics when fish treated with free Dex 24 h after induction of rod cell ablation with Mtz were compared to Mtz-only controls (9). Here, we used the same post-ablation treatment paradigm to compare Dex and D-Dex effects on rod cell regeneration kinetics (Fig. 3A). YFP-NTR expressing transgenic larvae were first exposed to 10 mM Mtz for 24 h from 5-6 dpf to induce rod cell ablation. Loss of YFP-expressing rod cells was visually confirmed using stereo-fluorescence microscopy. Mtz was then removed and larvae were either: i) returned to normal conditions (Mtz-only controls); ii) soaked in Dex (control for enhanced regeneration); or injected with iii) free Dex, or iv) D-Dex. At 9 dpf (72 h into the regenerative phase) YFP^+^ rod cell regeneration kinetics were measured using a fluorescent plate reader. Note, at this time point, roughly 50% of rod cells have regenerated in larvae with Mtz alone relative to non-ablated controls (23).

**Figure 3.**
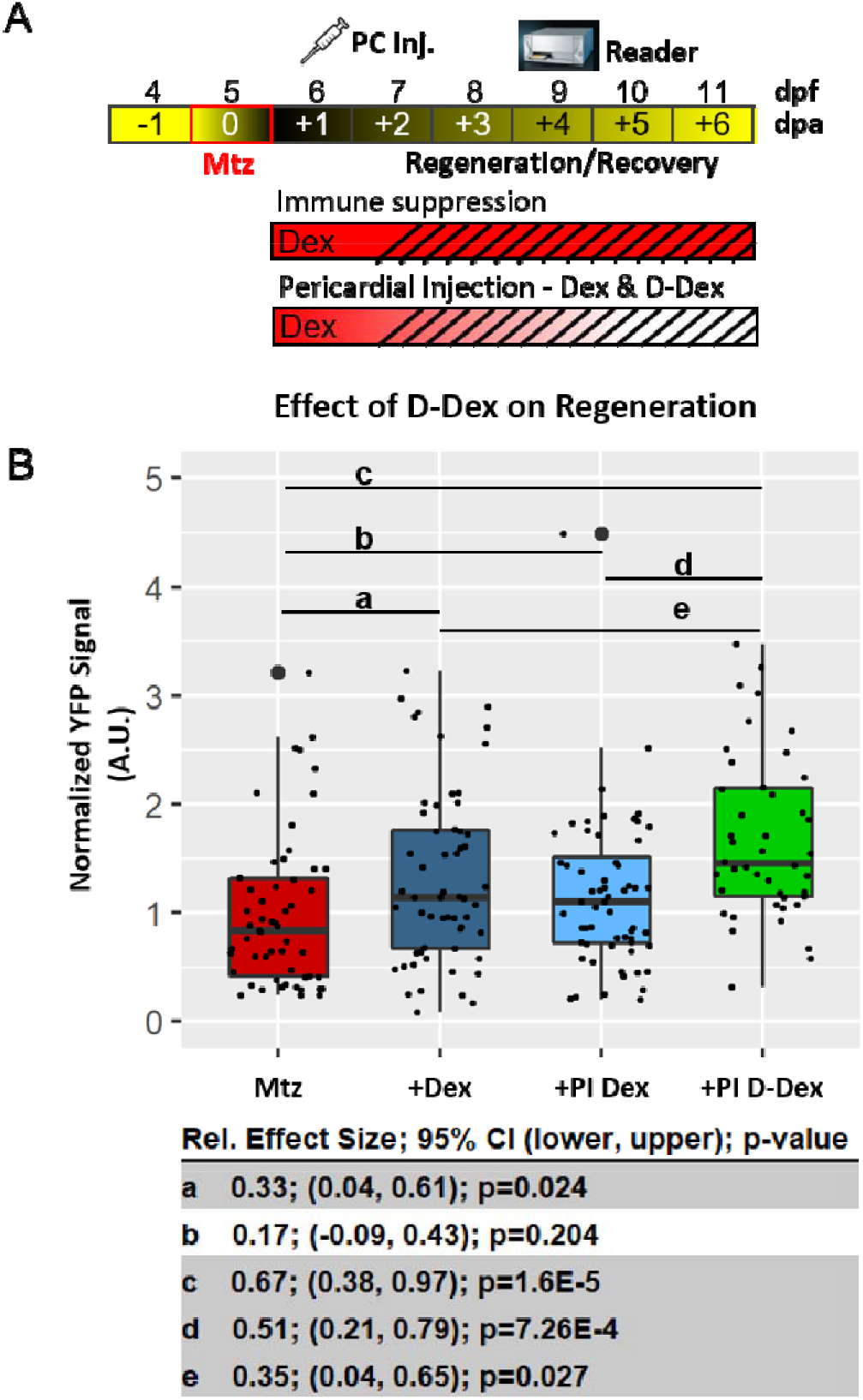
D-Dex treatment super-accelerates retinal regeneration compared to free Dex. **A,** Regeneration assay schematic, at 5 dpf fish expressing NTR-YFP in rod cells were separated into 4 groups, 1) ablated with Mtz (10 mM, 24 hours), and 2-4) ablated and subsequently treated with 2.5 μM Free-Dex (+Dex) or a pericardial injection (PI) of 5 μM Free-Dex (+PI Dex) or D-Dex (+PI D-Dex) at 6 dpf. Three days later (9 dpf), YFP labeled rods in all fish were quantified with a TECAN fluorescent plate reader. **B,** Boxplot of YFP signals across groups (Mtz only control, +Free-Dex, +PI Dex, +PI D-Dex) relative to control fish (Mtz only). Statistical comparisons between groups (a-e) were done to calculate effect size, 95% confidence intervals, and p-values between select conditions. +Dex enhanced rod cell regeneration by 33%, similar to our previous observations. +PI D-Dex further accelerated regeneration by 67% compared to control and 35% to +Dex only.

Compared to Mtz-only controls, all three Dex treated conditions resulted in enhanced rod cell regeneration kinetics (Fig. 3B). However, statistically significant increases were only observed for larvae soaked in Dex or injected with D-Dex. The increase in regeneration kinetics observed with soaking in Dex, ~30%, was identical to our previous findings (9). Intriguingly, larvae injected with D-Dex showed a ~67% enhancement in rod cell regeneration kinetics compared to Mtz-only treated controls (p=1.6E-05 D-Dex vs Mtz, n=45). This is nearly double the regeneration rate of larvae soaked in Dex (+34%, p=0.027 D-Dex vs Dex, n=58) (Fig. 3B).

### Dex does not promote rod cell survival

To control for the possibility that Dex treatment stimulates rod cell survival rather than promotes regeneration, i.e., acted as a neuroprotectant, we modified the protocol and measured YFP levels at 7 dpf instead of 9 dpf. At this time point, maximal loss of YFP signal is observed in Mtz-only controls (17), facilitating tests for neuroprotective effects, as per a large-scale screen we recently performed to identify compounds promoting rod cell survival (17). Here, compared to larvae treated with Mtz alone, treatment with Dex 24 h after induction of cell death had no effect on rod cell survival (Supp. Fig. 3).

### D-Dex localizes to dying rod photoreceptors and subsequently to reactive immune cells

We next assessed the dynamics of dendrimer cellular targeting using *in vivo* time series confocal imaging of Cy5-conjugated dendrimers (D-Cy5) after induction of rod cell death. Previous data from static imaging studies suggested dendrimer nanoparticles accumulate in reactive microglia/macrophages at sites of injury/inflammation (16). Here, time series imaging allowed changes in dendrimer localization to be assessed in individual fish over time (Fig. 4A, B). To monitor interactions between dendrimers, retinal microglia and rod cells, double transgenic fish expressing NTR-YFP in rods and tdTomato in microglia were used. To promote a sustained inflammatory environment, we prolonged the cell ablation process by treating larvae with a decreased concentration of Mtz (2.5 mM) for 48 h, from 5-7 dpf. Midway through this process, at 6 dpf, D-Cy5 was injected pericardially, followed immediately by time-series imaging. At 90 min post-injection, D-Cy5 co-localized with dying rod cells in the outer nuclear layer (ONL; Fig. 4C, inset). Although what are presumed to be reactive microglia were observed nearby, no quantitative evidence of co-localization with D-Cy5 and microglial cells was observed at this time. In contrast, at 270 min post-injection, tdTomato-expressing retinal microglia and peripheral macrophages co-localized strongly with D-Cy5 (Fig. 4D). To better monitor extended interactions between D-Cy5 and labeled cells, confocal time-lapse imaging was used as per our prior report (9). In a 4D-rendered video in which imaging begins 30 h post-Mtz treatment and 6 h post-injection of D-Cy5, frequent examples of microglia acquiring D-Cy5 are observed following initial D-Cy5 localization near dying rod cells (Supp. Video. 7).

**Figure 4.**
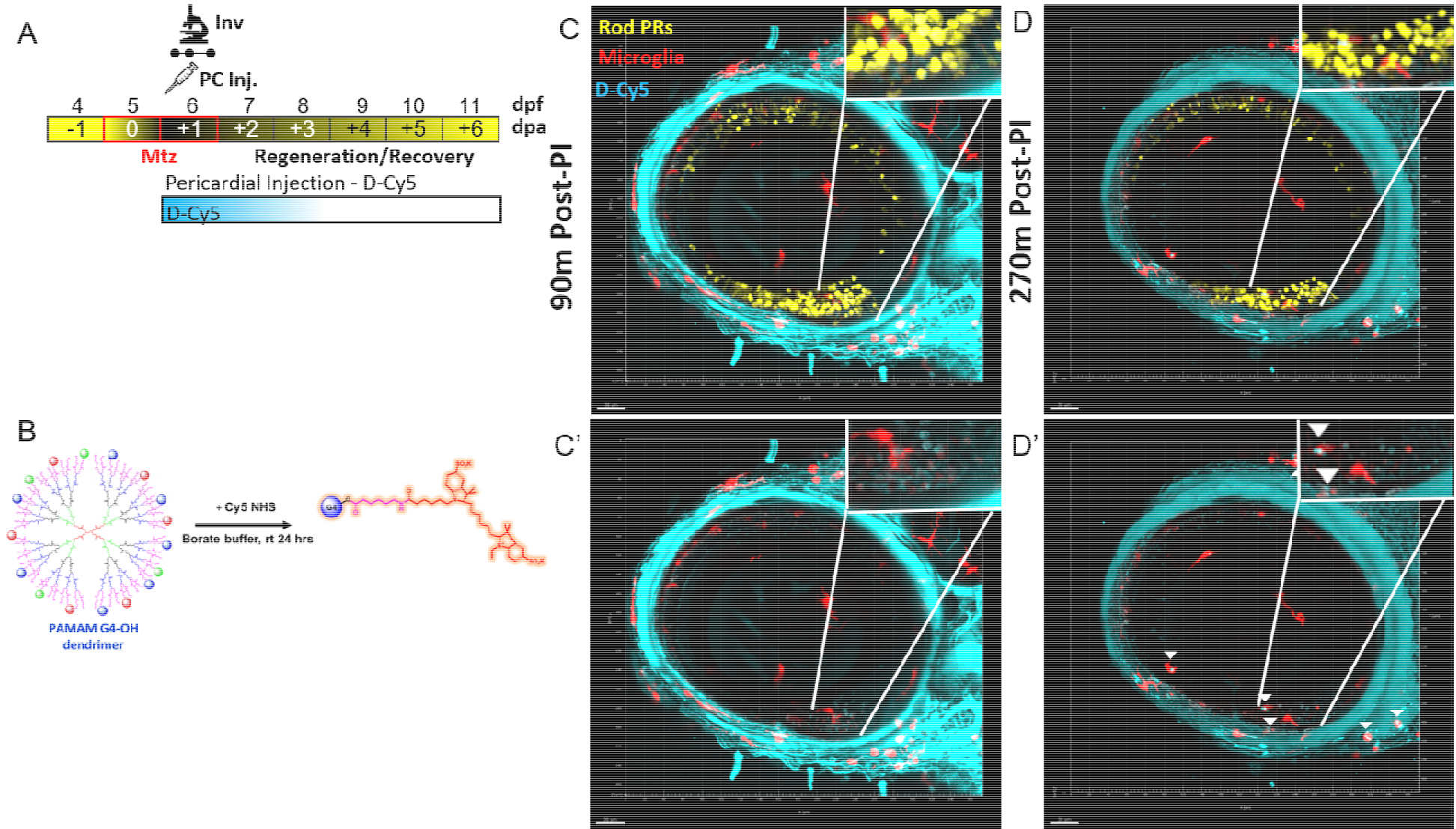
Dendrimer-Cy5 localizes to dying Rod Photoreceptors. **A,** Assay schematic, zebrafish expressing rod photoreceptors (yellow) and microglia/macrophages (red) were treated with 2.5 mM Mtz from 5-7 dpf to slow the cell death process. At 6 dpf, fish were given PC injections of dendrimers conjugated to Cy-5 fluorescent reporter (cyan, D-Cy5, 2 ng/ul) and immediately imaged using time series *in vivo* confocal microscopy. **B,** Creation of labeled D-Cy5 using PAMAM G4-OH, Cy5 NHS, and borate buffer. **C-D**, Image stills from *in vivo* confocal time-lapse microscopy. **C,** Image taken at 90 minutes post injection of D-Cy5, inset shows microglia interacting with rod cells. **C’**, Same image minus YFP channel, inset of same region as in C showing D-Cy5 presumably localized to dying rod cells in the outer nuclear layer (ONL) but not yet co-localized with microglia. **D,** Image taken at 270 minutes post injection of D-Cy5, inset shows microglia interacting with rod cells. **D’,** Same image minus YFP channel, inset show microglia (red) that have now taken up D-Cy5 (cyan, white arrowheads). See corresponding supplemental video 7.

## Discussion

We used AO-LLSM-based *in vivo* time-lapse imaging to investigate microglia/macrophage dynamics during retinal regeneration and leveraged dendrimer nanoparticles for targeted delivery of Dex to reactive macrophage/microglia. AO-LLSM-based time-lapse imaging revealed that post-ablation Dex treatment significantly decreased microglia migration speed but minimally altered microglia morphology; i.e., ameboid morphologies associated with reactivity were maintained. We interpret this as an immunosuppressive effect, despite sustained sphericity, linking inhibition of a specific microglia behavior to enhanced neuronal regeneration kinetics. Furthermore, we show that dendrimer-Dex formulations super-accelerate retinal regeneration kinetics in zebrafish, supporting their use in stimulating the regenerative potential of retinal MG and perhaps neuronal stem cells of the CNS more broadly.

The role of the immune system in neurodegenerative disease is complex, with evidence of both beneficial and deleterious roles (11, 24–26). Data from our lab and others support context-dependent roles for microglia during neural regeneration, with evidence of both pro- and anti-regenerative effects (9, 10, 27, 28). Collectively, these data support the concept that modulating macrophage/microglia reactivity may provide a therapeutic strategy for promoting neural repair in humans (29, 30). We previously implicated retinal microglia in the regulation of rod photoreceptor regeneration (9). Consistent with a biphasic role, the immunosuppressant Dex had opposing effects on rod cell replacement kinetics: pre-injury treatments were inhibitory but post-injury treatments accelerated regeneration. In the previous report, time-lapse *in vivo* imaging showed that pre-injury Dex exposure inhibited microglia reactivity to rod cell loss; the effects of pro-regenerative post-injury Dex treatments on microglia behavior were not investigated. In addition, we noted deleterious effects when Dex treatments were prolonged or administered at higher doses. Our new imaging data provided novel insight into the mechanism of immunosuppression on microglia reactivity, that is consistent with microglia transitioning from a pro-reparative to anti-reparative state during retinal regeneration. This transition has been observed in other neural injury and neurodegenerative paradigms (31), but stands in contrast to classical M1 to M2 transitions observed in non-CNS disease contexts (32). Follow-up investigations of microglia function will be required to determine: 1) if reducing migration speed alone is sufficient to enhance regeneration kinetics, 2) when transitions to homeostatic morphologies occur, and 3) how post-ablation Dex alters gene expression profiles of microglia and MG stem cells. Interestingly, we also noted a range of morphologies and behaviors among neighboring microglia in close proximity to dying rod cells (Supp. Video 3). Microglia exhibit structural and transcriptomic heterogeneity both spatially and when comparing healthy and diseased tissue (33, 34), suggesting functional divergence. Single-cell transcriptomic studies have revealed signatures of neurodegenerative disease-associated microglia (DAM) (35, 36) and bulk RNA studies have delineated regeneration-associated microglia (RAM; (37)). That functionally discrete microglia subtypes exist was recently confirmed in a mouse model of retinal degeneration (38) and in zebrafish brain (39). AO-LLSM imaging data suggests an extension of this concept, with functional divergence arising even among neighboring microglia. Collectively, these studies highlight the need for comprehensive interrogations of microglia/macrophage subtype function during neuronal degeneration and regeneration, as well as the value of nanoparticle-based strategies for targeting reactive immune cells.

Clinical trials employing systemic immunosuppression/anti-inflammatories as a strategy for ameliorating neurodegenerative disease have failed (40). Targeted delivery approaches are therefore needed to harness potential benefits of this powerful family of drugs. Nanoparticle-based targeted immunomodulation has transformative therapeutic potential in this regard (41), PAMAM dendrimers provide a unique opportunity to selectively target reactive immune cells in CNS disorders. Dendrimers have also been shown to reduce drug toxicity, presumably due to rapid clearance in the absence of inflammation (42). Reporter-conjugated dendrimers also enable assessments of immune cell dynamics and function in experimental models, including regenerative paradigms. In keeping with prior results, we hypothesized that dendrimer-based targeted drug delivery of Dex to reactive microglial would enhance the pro-regenerative effects of post-injury immunosuppression we observed previously (9). This hypothesis was proven true, as nanoparticle-conjugated Dex showed consistent improvements compared to Dex alone. Specifically, conjugating Dex to G4 PAMAM dendrimers (D-Dex): 1) reduced Dex toxicity, 2) facilitated neuronal and immune cell targeting - initially targeting dying neurons and subsequently accumulating in phagocytic immune cells, and 3) super-accelerated neuronal regeneration kinetics in the zebrafish retina. Each of these findings is novel and builds upon an expanding body of work demonstrating profound therapeutic potential of dendrimer nanoparticles (12, 14, 15). Broadly, nanoparticle-based modulation of immune system responses to neuronal cell death may represent a generalizable strategy for enhancing neural stem cell activity. However, accelerated regenerative processes in zebrafish do not necessarily equate to improved functional recovery (43, 44) nor to enhanced regenerative capacities in mammals. Additional studies will be required to test how accelerated retinal cell regeneration correlates to the recovery of visual deficits and to assess targeted immunomodulation in mammalian retinal degeneration models. These findings nevertheless add to a growing appreciation of the importance neuroimmune interactions in disease, underscoring an urgent need to expand our understanding of the roles played by discrete immune cell types in neurodegenerative and neuroregenerative paradigms, and to develop next-generation strategies for targeted immunomodulatory therapies.

## Materials and Methods

### Zebrafish husbandry and transgenic lines

All studies were carried out in accordance with recommendations by the Office of Laboratory Animal Welfare (OLAW) for zebrafish studies and an approved Johns Hopkins University Animal Care and Use Committee animal protocol. All fish were maintained using established conditions at 28.5◦C with a 14:10 h light:dark cycle. Previously published transgenic lines used here included: *Tg(rho:YFP Eco. NfsB)gmc500* (23), *Tg(mfap4.1:Tomato-CAAX)xt6* (45) and *Tg(mpeg1.1:LOX2272-LOXP-tdTomato-LOX2272-Cerulean-LOXP-EYFP)w201* (46). *Tg(rho:YFP Eco. NfsB)gmc500* expresses a bacterial Nitroreductase (NTR) enzyme selectively in rod cells to enable selective ablation (see below). This line was further propagated in a pigmentation mutant, roy^a9^ (roy), to facilitate detection of YFP signal *in vivo*.

### NTR-Mtz mediated rod photoreceptor ablation and free-Dex treatment

NTR:YFP expressing larvae were separated into equal-sized groups at 5 dpf: (i) nontreated controls and (ii) larvae treated with either 10 mM Metronidazole (Mtz, Acros) for 24 h or 2.5 mM Mtz for 48 h at 5 dpf (specifically to slow down the injury process for *in vivo* confocal imaging). Mtz is reduced by NTR into a cytotoxic substance inducing DNA-damage followed by specific cell death. After Mtz treatment, fish were rinsed and kept in 0.3Å~ Danieau’s solution until quantified, imaged or killed. For experiments with non-conjugated Dex treatment, equal size groups of fish ablated with Mtz were treated with 2.5 μM Free-Dex. Fish treated with Dex only as controls were additionally used as well as untreated fish.

### Adaptive optics lattice light-sheet microscopy (AO-LLSM)

5-7 dpf zebrafish were anesthetized using tricaine (0.16 mg/ml) and then embedded in a 0.8% low melt agarose droplet on a 25-mm coverslip. A homemade hair-loop was used to position and orient fish. The excitation and detection objectives along with the 25-mm coverslip were immersed in ~40 ml of E3 media + PTU at room temperature (22 ± 1° C). Zebrafish expressing *Tg(rho:YFP Eco. NfsB)gmc500* and *Tg(mfap4.1:Tomato-CAAX)xt6* to label rod photoreceptors and microglia/macrophages, respectively (9)—were excited using 514 nm and 560 nm lasers simultaneously (514 nm operating at ~3 mW and 560 nm operating at 5 mW corresponding to ~15 μW and ~25 μW at the back aperture of the excitation objective) with an exposure time of 20-50 msec. Dithering lattice light-sheet patterns with an inner/outer numerical aperture of 0.36/0.4 for both colors were used. The optical sections were collected by scanning the sample stage with 400-500 nm step size, equivalent to 200-250 nm axial step size in detection objective coordinate, with a total of 121-201 steps. Emission light from the two fluorophores were separated by a long-pass filter (Di03-R561-t3, Semrock) and captured by two Hamamatsu ORCA-Flash 4.0 sCMOS cameras (Hamamatsu Photonics). Prior to the acquisition of the time series data consisting of 100–200 time points, the imaged volume was corrected for optical aberrations using two-photon “guide star” based adaptive optics method (manuscript in preparation). Each imaged volume was deskewed and deconvolved with experimentally measured point spread functions obtained from 100 nm tetraspec beads (Thermo Fisher) in C++ using Richardson-Lucy algorithm on HHMI Janelia Research Campus’ computing cluster. The AO-LLSM was operated using a custom LabVIEW software (National Instruments). Image analysis was performed using FiJi (i.e., ImageJ v1.52p; NIH) or Imaris (v9.5.1; Bitplane) to quantify by analysis including displacement and speed of microglia.

### Synthesis of fluorescently labeled dendrimer (D-Cy5, D-FITC) and dendrimer-dexamethasone (D-Dex) conjugates

To enable imaging of dendrimer in this zebrafish model, we fluorescently labeled the hydroxyl terminated generation 4 PAMAM dendrimers (G4-OH) by covalently conjugating a near IR dye Cyanine 5 (Cy5) to the surface -OH groups of the dendrimers to obtain D-Cy5. These protocols have been previously published (47, 48). The D-Cy5 conjugate was characterized for its loading and purity using proton-nuclear magnetic resonance (^1^H NMR) and high-performance liquid chromatography (HPLC) respectively. Similarly, Fluorescein isothiocyanate (FITC)-labeled dendrimer (D-FITC) was used for some biodistribution studies. D-FITC synthesis and characterization were performed as described previously (49). Dendrimer-dexamethasone conjugate (D-Dex) was synthesized by following previously reported synthesis procedure (22). Briefly, the D-Dex conjugates were synthesized using a two-step procedure. In the first step, dexamethasone-21-glutarate (Dex-linker) was synthesized by reacting the -OH group at the 21^st^ position of Dex to the -COOH of the glutaric acid in the presence of triethylamine as a base. The Dex-Linker was purified using flash column chromatography. In the second step, the Dex-Linker was conjugated to the -OH groups on the dendrimer surface in the presence of coupling agent benzotriazol-1-yloxy tripyrrolidinophosphonium hexafluorophosphate diisopropylethylamine (PyBOP) as a base and anhydrous dimethylformamide (DMF) as solvent. The final product was purified using dialysis against DMF to remove reacted Dex-linker and PyBOP side products for 24 hours followed by dialysis against water to remove DMF. D-Dex conjugate was characterized using ^1^H NMR to estimate the drug loading and HLPC for its purity.

### Intravital confocal microscopy

All intravital imaging applied previously detailed protocols (50). Fish expressing YFP and NTR in rod photoreceptors, *Tg(rho:YFP Eco. NfsB)gmc500*, and tdTomato in microglia/macrophages, *Tg(mpeg1.1:LOX2272-LOXP-tdTomato-LOX2272-Cerulean-LOXP-EYFP)w201*, were used. Confocal z-stacks encompassing the entire orbit of the eye or entire fish (step size, 5 microns) were collected at 10- or 20-min intervals over a total of between 4.5 and 20 hours. Image analysis was performed using FiJi (i.e., ImageJ v1.49b; NIH) or Imaris (v7.6.5; Bitplane) to quantify by morphometric analysis as sphericity, correlation between area and volume or by volume in the kidney using minimum bounding spheres.

### IMARIS volumetric rendering and quantification

Larval whole-retina images were collected as described above using identical acquisition and IMARIS processing parameters across all conditions. Photoreceptor volume was calculated using IMARIS local background-based volumetric rendering of YFP signals.

### ARQiv scans to measure rod photoreceptor regeneration kinetics

*Tg(rho:YFP Eco. NfsB)gmc500* larvae were treated with Mtz ± Dex, and regeneration kinetics were analyzed using the ARQiv system, as previously described (9). 5 dpf larvae were exposed to 10 mM Mtz for 24 hours and then were treated with a single dose of 2.5 μM Free-Dex or treated with D-Dex (see below). The ARQiv scan was performed on day 9.

### Pericardial injection of Dex and dendrimer-conjugated Dex and D-Cy5

*Tg(rho:YFP Eco. NfsB)gmc500* larvae were treated with Mtz at 5 dpf, and then at 6 dpf larvae were anesthetized with Tricaine, placed under a PLI-100 picospritzer (Harvard Apparatus) and injected with 10 nL amount of either 5 μM Dex, D-Dex or D-Cy5. The concentration was doubled compared to Free-Dex to account for differential uptake under injection compared to soaking.

Subsequently at 9 dpf, the ARQiv scan was performed to measure regeneration kinetics as above. In later experiments, the injection was performed with D-Cy5 in order to observe Dendrimer activity in *vivo*. Toxicity of this injection method was determined following injection of equal groups of fish with vehicle, and various concentrations of Free-Dex, or D-Dex (containing equivalent Dex in conjugated form) at 5 dpf. At 7 dpf, % of surviving fish was counted (Fig 2), LD_50_ was calculated using previously established methods (23).

### Statistical analysis

Data were processed with a custom R-based package (ggplot2) to generate box plots showing the first quartile (lower box), median (bold line), third quartile (upper box), upper and lower adjacent (whiskers), and raw data (dot plot; large dots denote outlier observations) for each experimental condition. Statistical analyses were carried out with R 3.3.1 and RStudio 0.99.893. Student’s t test was used to calculate effect size between paired groups with effect size, 95% confidence intervals, and P values provided.

## Supporting information

SuppVid4

SuppVid7

SuppVid2

SuppVid6

SuppVid5

SuppVid3

SuppVid1

SuppVidsDescriptions

## Acknowledgements

This work was supported by the following grants from the National Institutes of Health, R01OD020376 (JSM), R01EY025304 (RMK), and P30EY001765-45 (core grant to Wilmer Eye Institute). We thank the Ramakrishnan and Tobin laboratories for sharing transgenic fish, and members of the Mumm lab for providing helpful discussions.

---

**Supplemental Figure 1.**
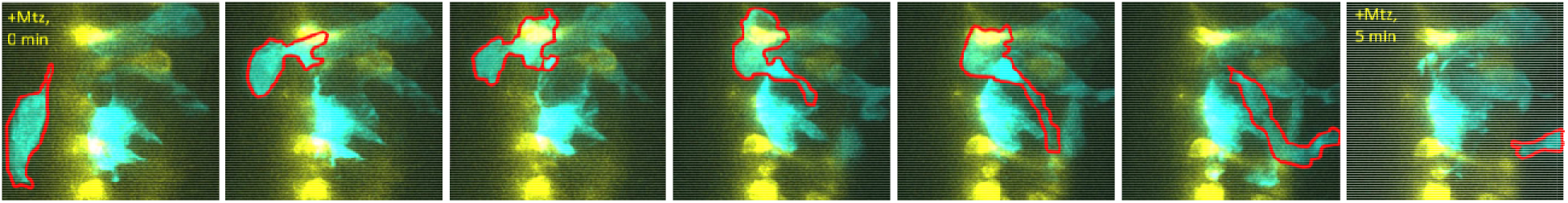
Peripheral macrophage translocation into the retina after induced rod cell death. Image stills from AO-LLSM time-lapse imaging of a single fish retina over a five-minute time period (left to right). Rod cells (yellow, NTR-YFP) were ablated with Mtz 12 hrs prior to imaging. A number of microglia/macrophages (cyan, false coloring of tdTomato) have aggregated in the outer nuclear layer near dying rod cells. A peripheral macrophage (outlined in red) appears to migrate from outside of the retina, or possibly from the retinal pigment epithelial layer, to inside the retina. See supplemental video 5 for corresponding video and supplemental video 6 for a second example.

**Supplemental Figure 2.**
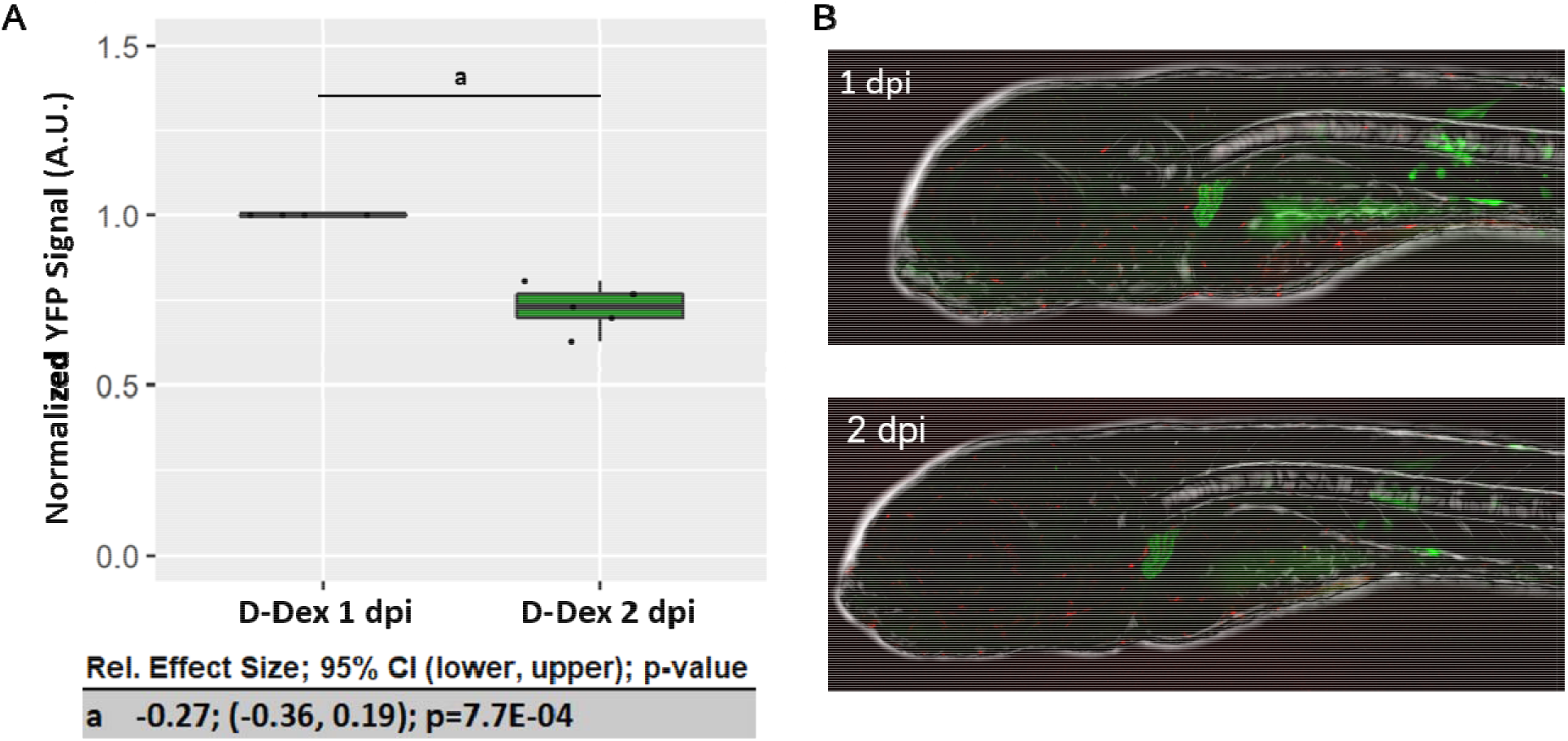
D-FITC accumulates in the kidney and is rapidly cleared. **(A-B)** To measure pharmacokinetic clearance rates of dendrimers, D-FITC (green) was injected into the pericardium of 5 mfap4:tomato-CAAX transgenic fish at 5 dpf(microglia/macrophages, red). **A,** Imaris was used to calculate the volume of D-FITC in the kidney at 6 and 7 dpf (1 and 2 dpi, days post injection). Signal at 2 dpi was normalized to 1 dpi. Statistical comparisons were done to calculate effect size, 95% confidence intervals, and p-value. A 27% reduction in D-FITC was seen between 1 and 2 dpi, suggestive of effective and rapid clearance in larval zebrafish. In no cases was accumulation of D-FITC observed in unperturbed retinas. **B,** Representative images for one fish at 1 and 2 dpi.

**Supplemental Figure 3.**
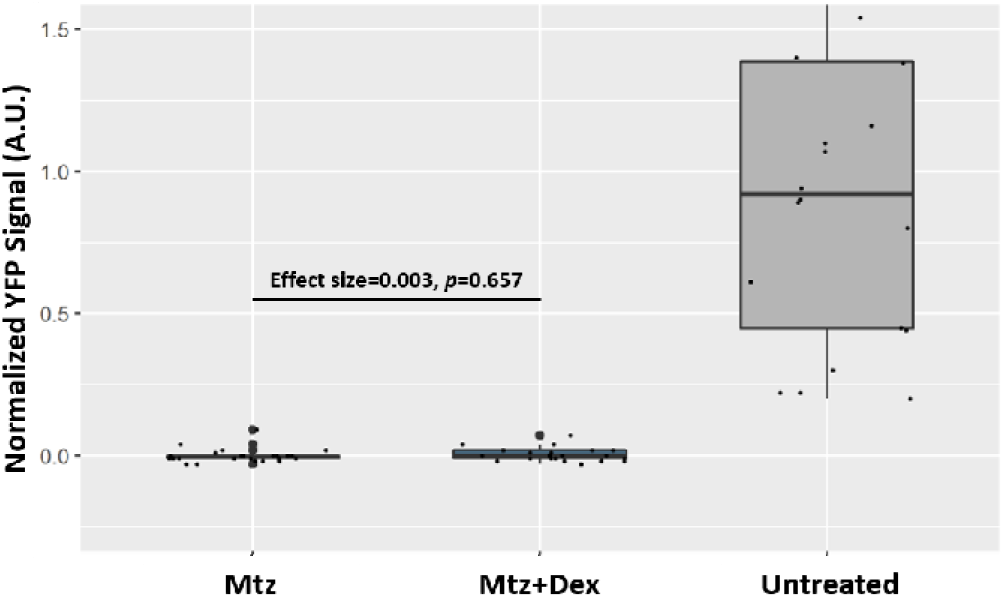
Dex is not neuroprotective of rod photoreceptor death. To test if Dex treatments protected NTR-YFP expressing rod cells from Mtz-mediated cell death, fish were treated with 2.5 μM Free-Dex (+Dex) 4 hours prior and throughout induction of rod cell ablation (10 mM Mtz, 24 hours, 5-6 dpf). At 7 dpf, YFP levels were quantified using an established plate reader assay. YFP signals are plotted relative to unablated controls (Untreated, set at 1.00) and ablated controls (Mtz only, set at 0) and statistical comparisons between the Mtz and Mtz+Dex groups were done to calculate effect size, 95% confidence intervals, and p-value. The addition of Dex had no effect on the rate of rod cell death.

