## Supplementary material for "Dendrimer-targeted immunosuppression of microglia reactivity super-accelerates photoreceptor regeneration kinetics in the zebrafish retina": SuppVidsDescriptions

Supplemental Video 1. Microglia response in control fish

Representative time-lapse video for untreated control fish expressing Rod PRs (yellow, NTR-YFP) and microglia (cyan, false coloring of tdTomato). Imaging was done using AO-LLSM.

Supplemental Video 2. Microglia response in fish treated with Dex only

Representative time-lapse video for Dex-only treated (2.5 µM, 24 hours) fish expressing Rod PRs (yellow, NTR-YFP) and microglia (cyan, false coloring of tdTomato). Imaging was done using AO-LLSM.

Supplemental Video 3. Microglia response in Mtz ablated fish

Representative time-lapse video for fish expressing Rod PRs (yellow, NTR-YFP) and microglia (cyan, false coloring of tdTomato) following treatment with Mtz (10 mM, 12 hours). Imaging was done using AO-LLSM.

Supplemental Video 4. Microglia response in Mtz and Dex treated fish

Representative time-lapse video for fish expressing Rod PRs (yellow, NTR-YFP) and microglia (cyan, false coloring of tdTomato) following treatment with Mtz (10 mM, 12 hours) and Dex (2.5 µM, 24 hours). Imaging was done using AO-LLSM.

Supplemental Video 5. Apparent microglia translocation inside the retina

Time-lapse imaging using AO-LLSM over a five-minute time period of rod PRs (yellow, NTR-YFP) ablated with MTZ (10 mM, 12 hours) in a fish co-labeled for microglia (cyan, false coloring of tdTomato). Shows what appears to be the translocation of peripheral macrophage translocating from outside of the retina to inside the retina, or possible from the retinal pigment epithelium to inside of the outer nuclear layer.

Supplemental Video 6. Apparent microglia translocation inside the retina, second example

Time-lapse imaging using AO-LLSM over a five-minute time period of rod PRs (yellow, NTR-YFP) ablated with MTZ (10 mM, 12 hours) in a fish co-labeled for microglia (cyan, false coloring of tdTomato). Shows what appears to be the translocation of peripheral macrophage translocating from outside of the retina to inside the retina, or possible from the retinal pigment epithelium to inside of the outer nuclear layer.

Supplemental Video 7. 4D imaging of D-Cy5, Rod PRs and microglia

Time lapse video of co-labeled Rod PRs (YFP+), microglia (RFP+) and PI D-Cy5 (Cyan). Following Mtz treatment (5 dpf, 2.5 mM, 48 hours) and D-Cy5 injection (6 dpf, 2ng/µL; 24 hours post-Mtz treatment), confocal imaging began six hours later (30 hours post-Mtz treatment). As Rod PRs are dying, D-Cy5 initially clusters near them. Subsequently microglia appear to phagocytose Rod PRs and acquire the D-Cy5 in several instances. For clarity, the video is repeated a second time without the Rod PR channel to easier visualize D-Cy5 and microglia interactions.
